# Overcoming the challenges of cascade reactions and complex substrates in natural product biocatalysis: Immobilization of a cyclodipeptide synthase

**DOI:** 10.1101/2024.07.24.604810

**Authors:** Lynette Alvarado-Ramírez, Emmajay Sutherland, Elda M. Melchor-Martínez, Roberto Parra-Saldívar, Alfredo D. Bonaccorso, Clarissa Melo Czekster

## Abstract

Cyclodipeptide synthases (CDPSs) use aminoacylated tRNAs to produce cyclic dipeptide natural products which can have anticancer and neuroprotective activity. Despite their potential, applications involving CDPSs are hindered by enzyme instability and challenges in producing aminoacylated tRNAs. Immobilizing enzymes can enhance stability and recyclability, yet studies on immobilized enzymes using aminoacylated tRNAs are lacking. Here, we immobilized the CDPS enzyme from *Parcubacteria bacterium* RAAC4_OD1_1 (PbCDPS) using three sustainable supports: biochar from waste materials, calcium-alginate beads, and chitosan beads. Active PbCDPS immobilization led to production of the cyclodipeptide cyclo (His-Glu) (cHE). Notably, following activation with glutaraldehyde, a five-fold increase in cHE production was observed, while the immobilized enzyme remained active for seven consecutive cycles. Furthermore, we co-immobilized three enzymes required for the cascade reaction yielding cHE, all of which require aminoacyl-tRNA substrates (PbCDPS, histidyl-tRNA synthetase, and glutamyl-tRNA synthetase). This enzymatic cascade successfully generated the cyclic dipeptide of interest, showcasing the potential of immobilizing complex enzymes operating in cascade on a single support. We demonstrated that tRNAs remained free in solution without adsorption onto beads. This work paves the way for the immobilization of enzymes utilizing tRNAs and potentially other complex substrates, expanding the spectrum of reactions exploitable with this technology.

**Graphical Abstract:**

- For Table of Contents Only

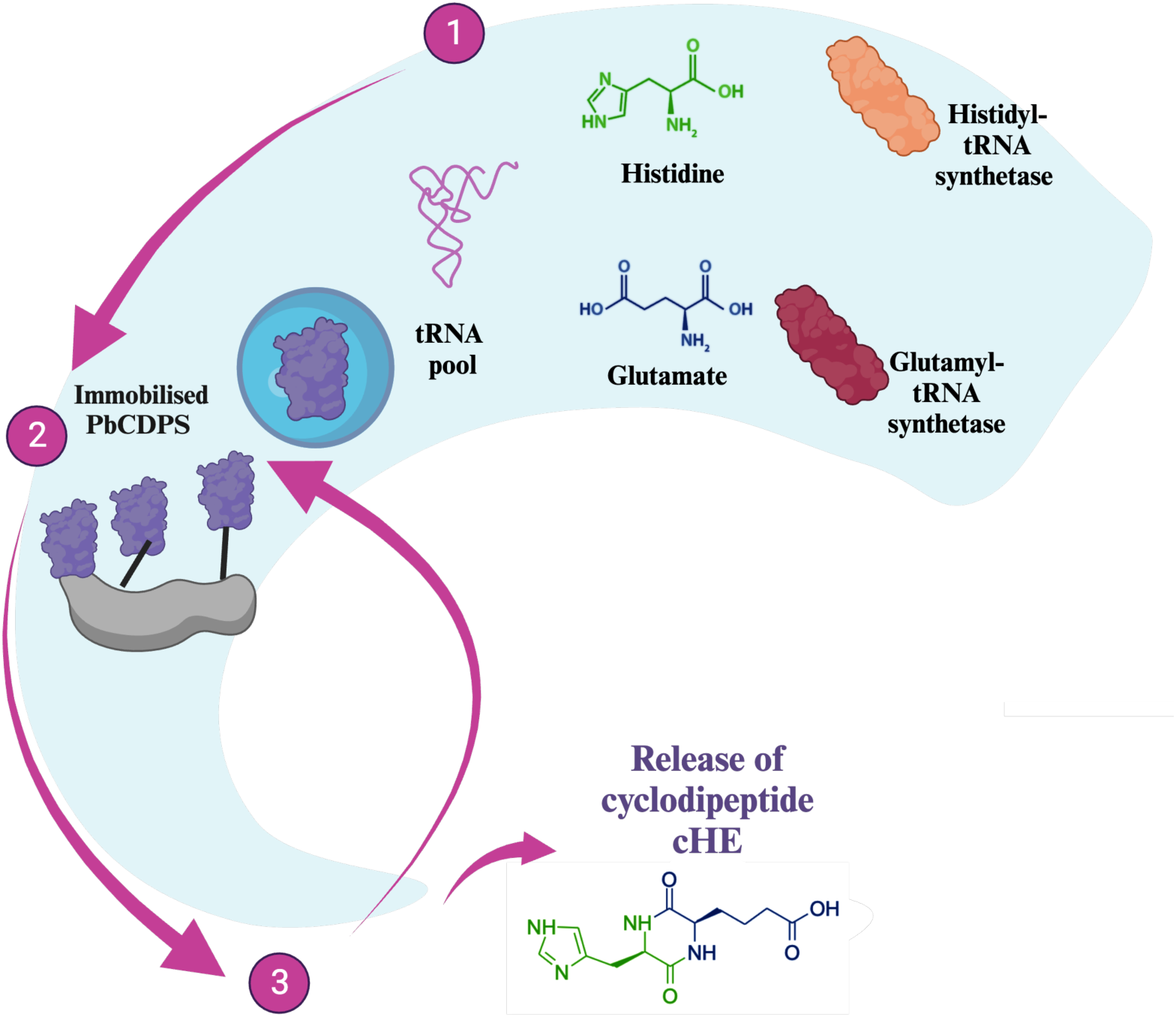

## Introduction

Cyclic dipeptides (CDPs) are natural products produced by organisms in all domains of life^1^. Multiple reports associate these compounds with anticancer activity^2–4^ and neuroprotection against neurodegenerative diseases such as amyotrophic lateral sclerosis^5^, Alzheimer’s^6^ and Parkinson’s disease^7^. CDPSs use two aminoacylated tRNAs (aa-tRNA) as substrates, and therefore rely on the activity of aminoacyl-tRNA synthetases (aaRS), hijacking aa-tRNAs from central metabolism and protein synthesis^9^. CDPS enzymes can be promiscuous, accepting multiple different substrates^10,11^, including unnatural amino acids^12,13^. Despite promising advances in the engineering of CDPSs^14^ expanding substrate scope, their usage in biocatalysis is limited due to cost, substrate complexity, and lack of recyclability^15,16^. The immobilization of enzymes participating in natural product biosynthesis has been limited to very few examples, including fluorinase^17^ and tyrosine decarboxylase^18^. Prior to this work, there were no reports of CDPS immobilization.

Immobilization allows the reutilization of enzyme catalysts, resulting in the reduction of operational cost^19,20^. Furthermore, enzyme immobilization offers several green chemistry advantages, such as enhanced stability and longevity, reduced environmental impact using sustainable supports, recyclability and reusability, and the utilization of waste-derived supports.^21^ When a support is used, the characteristics of the material can affect the immobilization process, and the utilization of inexpensive and sustainable support matrices could increase applicability to several enzymes^22^. Alginate and chitosan are natural biopolymers that serves as an excellent support for enzyme immobilization. Their attributes include being non-toxic, biodegradable, cost-effective, and capable of interacting with proteins due to the presence of hydroxyl and amino functional groups ^23–25^. Carbon materials are also suitable supports for enzyme immobilization ^26^. Biochar is a carbon-rich solid derived from the thermal decomposition of biomass (i.e., pyrolysis, flash carbonization, hydrothermal decomposition among others) from agro-industrial residues (coffee, algae, and others) ^27^. Biochar can be easily engineered, and the process is economically sustainable ^28^. With each support, different approaches can be used including entrapment, adsorption, and covalent bonding, and therefore careful optimization must be carried out.

Previously, the production of cyclic dipeptides with high yield using the CDPS enzyme from *Parcubacteria bacterium RAAC4_OD1_1* (PbCDPS, GenBank: ETB63777.1) was demonstrated^29^. We focused on the production of cyclo(His-Glu) (cHE), the major product from the PbCDPS-catalysed reaction, and currently not commercially available. PbCDPS can also produce cyclo(His-Pro) (cHP) as an additional reaction product^13^. Developing simple enzymatic routes to histidine-containing cyclodipeptides is an important step for the exploitation of these molecules as potential bioactive agents^30,31^. The CDPS reaction occurs in a cascade starting with amino acids histidine (His) and glutamate (Glu), including aaRS enzymes and ATP. It is therefore an ideal system to demonstrate the feasibility of including immobilization and catalyst recycling in a complex system involving aa-tRNA and three distinct enzymes.

Here, we describe a facile strategy for the immobilization of CDPS enzymes using three different supports: Ca-alginate, chitosan beads and biochar derived from various sources (macroalgae, spent coffee, distillery waste, polystyrene, and wood waste). We focused on the immobilization of PbCDPS, followed by histidyl tRNA synthetase (HisRS) and glutamyl tRNA synthetase (GluRS). The reusability of the system and the effects of quenching and loading in the support matrix were determined. We successfully immobilized PbCDPS alone and together with the complete enzymatic cascade for cHE production (PbCDPS, HisRS and GluRS). This work sets the stage for future studies employing enzyme immobilization of CDPSs and other enzymes that utilize tRNAs as substrates, enabling complex biocatalytic cascade reactions leading to the biosynthesis of natural products and analogues.

## Materials and methods

### Materials

4-(2-Hydroxyethyl)-1-Piperazineethanesulfonic acid (HEPES 99%) and potassium hydroxide were purchased from Fisher bioreagents. Potassium chloride (CaCl_2_), adenosine 5’-triphosphate disodium salt hydrate (ATP), sodium alginate, chitosan (low molecular weight), 2-mercaptoethanol (>99%) and imidazole were purchased from Sigma Aldrich. Magnesium chloride (MgCl_2_), sodium chloride (NaCl) and glutamic acid (Glu, 99%) were from Acros Organics. Histidine (His) free base was from from MP Biomedicals. Dithiothreitol (DTT), chloroform, trifluoracetic acid, LC-MS-grade and HPLC-grade water and acetonitrile were from Thermo Fisher. Isopropylthio-β-galactoside (ITPG, >99%) was from Neo Biotech. Glycerol and acid phenol:chloroform (>99%) were from Ambion. DEPC-treated water was prepared in house by adding diethylpyrocarbonate (Fisher Scientific) to a final concentration of 0.1 %, followed by stirring overnight followed by autoclave sterilization.

### PbCDPS and aaRS production and purification

PbCDPS was produced and purified following the protocol of Sutherland et al.^12^. The gene encoding PbCDPS was cloned into a pJ411 expression vector with a C-terminal hexahistidine tag and transformed into *E. coli* BL21(DE3) competent cells (NEB). Cells were grown at 37 °C until the OD_600_ reached 0.6 and protein expression was induced with IPTG (1mM). Cells were then grown at 16 °C overnight. After harvesting, the resultant cell pellet was resuspended in 30 mL per 1 L of grown culture of lysis buffer (50 mM HEPES pH 7.0, 250 mM NaCl, 20 mM imidazole, 5% glycerol). Cells were lysed using a high-pressure cell disruptor (Constant Systems), and centrifuged at 51000 g for 30 minutes, 4 °C. The lysate was filtered through an 0.8 µm membrane and loaded onto a 5 mL HisTrap HP column (GE Healthcare), pre-equilibrated with lysis buffer. The column was washed with 20 column volumes (CV) of lysis buffer and the adsorbed proteins were eluted using elution buffer (50 mM HEPES pH 7.0, 250 mM NaCl, 300 mM imidazole, 5% glycerol) with stepped increasing concentrations of imidazole (10%, 20% and 100%, 10 column volumes each). Proteins of interest were dialyzed into dialysis buffer (20 mM HEPES pH 7, 250 mM NaCl, 5 mM 2-mercaptoethanol) overnight at 4 °C. PbCDPS was further purified via size exclusion chromatography using a Superdex 200 Increase 16/60 column pre-equilibrated with dialysis buffer. Fractions containing pure PbCDPS were pooled, concentrated to 10 mg/mL, flash frozen in aliquots and kept at −80 °C for future use. Enzyme concentration was measured using the Nanodrop DeNovix (DS-11 FX) spectrophotometer/fluorometer.

The aminoacyl-tRNA synthetases of interest to this project – GluRS and HisRS – were purified from *E. coli* as detailed above following the protocol of Sutherland et al. ^12^. The purification buffers for GluRS contained 50 mM HEPES pH 8, 500 mM NaCl and 20 or 300 mM imidazole whilst HisRS buffers contained 50 mM HEPES-KOH pH 7.6, 10 mM MgCl_2_, 2 mM 2-mercaptoethanol and 10 or 400 mM imidazole. aaRS enzymes were concentrated to 10 mg/mL, flash frozen in aliquots and kept at −80 °C for future use.

### tRNA Pool Extraction

For the extraction of the pool of all tRNAs produced by *E. coli,* the protocol described by Sutherland et al^12^ was used without further modification.

### Synthesis of biochar

The biochar utilized in this study was prepared from various waste sources by using a conventional low-temperature pyrolysis methodology. These sources included: I) spent coffee obtained from a local coffee shop in St. Andrews, Scotland; II) a mixture of polystyrene & wood, composed of 95% wood and 5% polystyrene waste; III) dry distillery waste; IV) dried macroalgae sourced from the North Sea (Guardbridge, St. Andrews, Scotland) (Fig. 1.1).

**Figure 1.**
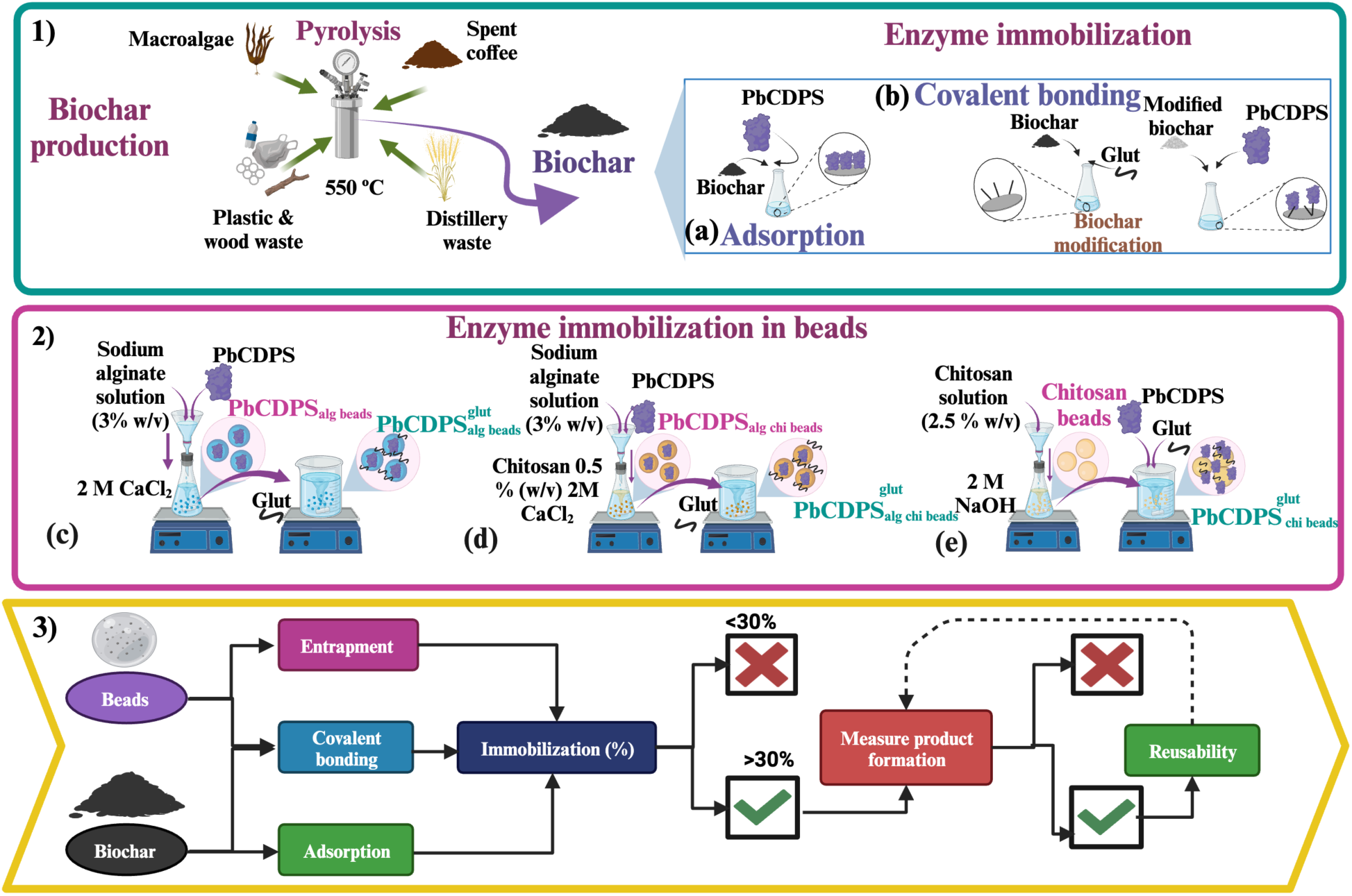
Workflow describing the evaluation and prioritization of different immobilization strategies. 1) Biochar production from spent coffee, macroalgae, polystyrene & wood and distillery waste, (a) PbCDPS immobilization on biochar through adsorption and (b) covalent bonding. 2) Process for PbCDPS immobilization in beads: (c) Ca-alginate beads, (d) alginate coated-chitosan beads, (e) chitosan beads. 3) Prioritization of the different immobilization strategies used for PbCDPS, starting with two different types of supports, the immobilization was made through entrapment, adsorption, and covalent bonding. Supports with more than 30% of immobilization were selected for the measurement of product formation (cHE) and the reusability was tested only for the supports with the higher concentration of the cHE.

Production of biochar was conducted using a pyrolysis reactor designed in-house. The reactor was a vertical-tubular design constructed with Stainless Steel 316 material. To ensure temperature homogeneity inside the reactor, the waste material was placed on an SS316 disc mesh that allowed the gas to cross the samples at the center of the reactor. Temperature monitoring was achieved using an internal thermocouple. Prior to initiating the pyrolysis process, nitrogen gas was purged into the reactor to eliminate any trace of oxygen. The waste products were subjected to pyrolysis at a temperature of 550°C, with a ramp rate of 10°C min^-1^ and a residence time of 30 minutes under a nitrogen flow of 50 mL min^-1^. Following the 30-minute duration at 550°C, the samples were rapidly cooled to prevent alterations in the residence time. Subsequently, the pristine biochar derived from the different waste sources underwent physical modification using various methods. This involved subjecting the biochar to thermal treatment at high temperatures in the presence of oxidizing agents such as carbon dioxide or steam. Additionally, depending on the specific biochar source, acid or hydrogen peroxide pre-treatment was employed.

For the biochar derived from distillery waste, polystyrene-wood waste, and coffee waste, a quartz tube was used to contain the material within the furnace. Two quartz-wool plugs were placed on either end of the tube to maintain the material at the center. The biochar was heated under a nitrogen atmosphere until it reached 920 °C. At this temperature, the gas atmosphere was switched from nitrogen to carbon dioxide. The physical modification process was carried out at 920 °C for 1 hour with a flow rate of 40 L min^-1^ of carbon dioxide. This treatment was used to modify the structure and porosity of the biochar, resulting in increased pore dimensions and surface area as well as Carbon: Oxygen ratio in the final biochar due to further thermal treatment a 920 °C under N_2_. As for the biochar obtained from macroalgae, a different approach was necessary due to its fine powder form. At 920°C, carbon dioxide was capable of completely gasifying the biochar. Therefore, hydrogen peroxide (H_2_O_2_) was employed as an oxidizing agent. The biochar was subjected to a temperature of 150 °C, which is the boiling point of hydrogen peroxide, for 1 hour in a reflux reactor. For the immobilization of PbCDPS on biochar, all sources were used with and without activation.

### PbCDPS immobilization

All experiments were performed in triplicate, data are reported as average ± SEM.

#### Biochar

##### Simple adsorption

A preliminary experiment to select the time of immobilization was carried out, analyzing the protein remaining in the supernatant following immobilization. Based on this, optimal time for immobilization was 30 minutes for adsorption methods. For the immobilization by physical adsorption of PbCDPS on biochar, 15 mg of biochar was mixed at room temperature, by gently stirring with 15 μg of PbCDPS in the presence of buffer (100 mM HEPES, 100 mM KCl and 10 mM MgCl_2_ at pH 7). After adsorption, immobilized PbCDPS was separated from the solution by centrifugation and washed with the same buffer to remove unattached protein from the support. Protein in the supernatant and buffer after washing were pooled and quantified to evaluate immobilization efficiency (in % PbCDPS immobilized).

##### Adsorption on activated biochar

To increase the number of functional groups in the support surface and allow a covalent link between the enzyme and the support, biochar was impregnated with glutaraldehyde ^32^. The impregnation was made by mixing the support with a glutaraldehyde solution (2% v/v) for 2 h at room temperature. Subsequently, biochar was washed with deionized water to eliminate excess glutaraldehyde. Following this, 15 mg of modified biochar was mixed with 100 µg of PbCDPS for 30 min at room temperature. After immobilization, the support was washed (100 mM HEPES, 100 mM KCl and 10 mM MgCl_2_ at pH 7) to remove excess protein. Protein in the supernatant and buffer after washing were pooled and quantified to evaluate immobilization percentage. Immobilization percentage of PbCPDS on biochar was determined from the difference between the initial protein and protein detected in the supernatant and washes. The total protein concentration was assessed using the Bradford method at a wavelength of 595 nm.

#### Alginate or chitosan beads

##### Alginate beads

PbCDPS was entrapped in alginate beads by mixing 115 mg of enzyme with 1 mL of sodium alginate solution (3 % w/v), following a reported protocol ^33^. The mixture was stirred and dropped through a syringe into 10 mL of CaCl_2_ 0.2 M. After, *PbCDPS_alg beads_* were washed with a HEPES/KCl/MgCl_2_ at pH 7 buffer to remove free enzyme. The CaCl_2_ and solutions from washes were collected to evaluate immobilization percentage and efficiency. *PbCDPS_alg beads_* were activated with 2% v/v glutaraldehyde for 2h at 4 °C, to enhance the number of functional groups on the support surface and enable covalent bonding between the enzyme and the support. Subsequently these beads are referred to as 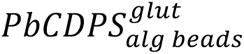.

##### Alginate-coated chitosan beads

Alginate-coated chitosan beads were prepared according to the optimal conditions described by a reported protocol ^34^. A solution containing a 3% w/v solution of sodium alginate and 115 mg of PbCDPS were added as drops using a syringe into a solution containing 10 mL of CaCl_2_ 0.2 M, 0.2% chitosan and 1.5% of acetic acid pH 5. Subsequently, *PbCDPS_alg:chi beads_* were washed with buffer (100 mM HEPES, 100 mM KCl and 10 mM MgCl_2_ at pH 7) to remove free enzyme. The chitosan/acetic acid/CaCl_2_ solution and washes were collected to evaluate immobilization percentage and efficiency. *PbCDPS_alg:chi beads_* were further activated with 2% v/v glutaraldehyde for 2 h at 4 °C, and these beads were subsequently referred to as 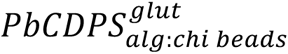.

##### Chitosan beads

Chitosan beads were prepared according to a protocol previously described ^35^ with some modifications. A solution of 2.5 % w/v of chitosan and 1.5% acetic acid was prepared and extruded dropwise into 2M NaOH solution at room temperature. After, chitosan beads were collected and washed with deionized water to remove excess NaOH. Following this, chitosan beads were activated using 2% v/v glutaraldehyde for 2 h at room temperature. Excess glutaraldehyde was removed by washing the activated beads with deionized water. Finally, beads were incubated with PbCDPS overnight at 4 °C for immobilization. Activated chitosan beads 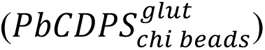 were washed with buffer (100 mM HEPES, 100 mM KCl and 10 mM MgCl_2_ at pH 7) to eliminate free enzymes. All buffers and solutions from washes were collected to evaluate immobilization percentage and efficiency.

### Characterization of biochar and beads

The functional groups of biochar and beads were evaluated using Fourier transform infrared technique (FTIR: Shimadzu IR Affinity 1S IR Spectrometer – for solid or liquid samples). Specific surface area (Brunauer-Emmett-Teller, BET) and pore size and volume (BJH) were determined using a Micromeritics Tristar ii Surface Area and Porosity Instrument with VacPrep 061 Degasser by adsorption and desorption of nitrogen at 77 K. The morphology was analyzed using a scanning electron microscopy (SEM) EVO MA 25 ZEISS.

### Determination of enzyme activity and product formation cHE

All reactions to determine product formation were performed in triplicate, data are reported as average ± SEM.

Production of cyclic dipeptide product by PbCDPS free in solution as well as after immobilization was detected using HPLC assays. Product identity was further verified by high resolution liquid chromatography mass spectrometry (LC-HRMS), following the protocol previously described by Sutherland et al^12^ for determination of cHE product without modification. Briefly, LC-HRMS was carried out using a Waters ACQUITY UPLC liquid chromatography system coupled to Xevo G2-XS QTof mass spectrometer equipped with an electrospray ionization (ESI) source. 10 µl of each sample were loaded onto an HSS-T3 column (2.1 × 100 mm, 1.8 µm, Waters Acquity, column temperature set to 40 °C). A gradient mobile phase ranging from 1 % B to 50 % B for 9 minutes was used, where the mobile phases consist of 0.1 % formic acid in water (A) and 0.1 % formic acid in acetonitrile (B) at a flow rate of 0.3 mL/min. The capillary voltage was set at 2.5 kV in positive ion mode. An MS^e^ scan was performed between 50 to 700 m/z. The m/z expected and observed for cHE were 267.1088 and 267.1093 respectively (ppm deviation: <2).

Reaction buffer contained 100 mM HEPES pH 7, 100 mM KCl, 10 mM MgCl_2_, 5 mM ATP, 10 mM DTT, 500 μM histidine, 500 μM of glutamate and 50 μM of tRNA pool. DEPC-treated water was added to achieve the final volume of 50 or 100 μl. Finally, 5 μM of each aaRS enzyme (HisRS and GluRS) and PbCDPS were added, and the reaction proceeded at least 16h at room temperature. Cold methanol was added to a final volume of 80% to quench the reaction. Samples were incubated at −80 °C for 15 min and centrifuged for 10 min at room temperature. The supernatant was transferred to a second Eppendorf tube and dried using nitrogen. Finally, LC-MS grade water was used to reconstitute samples to the same initial reaction volume.

The production of cHE was monitored using high performance liquid chromatography (HPLC, Shimadzu CMB-20A). Samples (20 μl) were injected onto a Waters XSelect Premier HSS-T3 column (4.6 x 50 mm, 2.5 mm) and run at 40 °C for 30 min. A gradient mobile phase from 1 % B to 50 % B over 5 minutes at a flow rate of 1 mL min^-1^ was used, where mobile phase A = 0.1 % trifluoroacetic acid in water and mobile phase B = 100 % acetonitrile. Absorbance was monitored at 214 nm and 254 nm.

A large-scale reaction (5 mL total volume) was set up to produce a cHE standard to be used for quantification. After the reaction proceeded for at least 16h, the mixture was separated using a 10 kDa filter membrane to remove enzymes and tRNAs present in the reaction. Then, the mixture was centrifuged to remove precipitate and the supernatant dried under nitrogen. The subsequent residue was resuspended in LC-MS grade water and this solution injected on the HPLC. A Shimadzu CMB-20A equipped with a fraction collector and a Waters XSelect Premier HSS-T3 column (4.6 x 50 mm, 2.5 mm) were used to purify cHE. The samples were injected and run at 40 °C for 30 min. A gradient mobile phase from 1 % B to 50 % B over 5 minutes at a flow rate of 1 mL min^-1^ was used, where mobile phase A = 0.1 % trifluoroacetic acid in water and mobile phase B = 100 % acetonitrile. The selected fractions were pooled and dried by lyophilization. Finally, this purified powder was used for LC-HRMS and as a standard for HPLC quantification.

### Optimization of loading and quenching during immobilization

All experiments were performed in triplicate, data are reported as average ± SEM. Different concentrations of protein were tested to determine the optimal amount of PbCDPS to be immobilized in each support (alginate and chitosan beads, biochar from spent coffee), as well as the effect in the production of cHE. For alginate beads 9.2, 13.9, 17.3 and 23.1 and μg PbCDPS were immobilized per mL of alginate. For biochar from spent coffee, 10, 15 and 20 μg of PbCDPS were used and finally for chitosan beads, 7.3, 14.6, 29.2 and 58.4 μg of PbCDPS were immobilized. The immobilization efficiency was measured as in the previous section, and the ratio of cHE produced with the immobilized PbCDPS per condition was compared. Additionally, to investigate if the support could retain or adsorb product after each reaction, we compared cHE obtained when the complete system was quenched (liquid and support) and when only the liquid was quenched.

### Reusability of immobilized PbCDPS

Reusability of immobilized PbCDPS was evaluated only with the supports that showed higher cHE production. After each overnight cycle (at least 16h reaction), supports under evaluation were recovered, and a new assay was started omitting PbCDPS but with other components as specified under “Determination of enzyme activity and product formation cHE”. All experiments were performed in triplicate, data are reported as average ± SEM.

### Co-immobilization of the enzymes involved in the cHE production

cHE is the final product of a cascade reaction between PbCDPS, HisRS and GluRS. Ideal conditions for PbCDPS were employed to immobilize the two other enzymes taking part in the cascade for cHE production. Simultaneous immobilization of PbCDPS, HisRS and GluRS was performed as described in the previous sections. Bradford was used to determine the amount of free protein and qualitatively estimate the immobilization efficiency. All experiments were performed in triplicate, data are reported as average ± SEM.

## Results and discussion

### Synthesis and characterization of biochar

Biochar produced from different sources by pyrolysis was used as carrier for the immobilization of PbCDPS. Biochar properties can be modified using an additional activation process after pyrolysis. Recent studies have shown that the activation with CO_2_ could increase specific surface area, pore structure and functional groups on the surface^28^. However, in previous experiments optimizing immobilization conditions (Table 1), high quantities of enzyme immobilized in the CO_2_-activated biochar resulted in undetectable cHE product formation. This could be due to the reduction of the pore size after the activation (Table S1), which limits diffusion of other reaction components such as aa-tRNA in and out of the biochar pores^36^.

**Table 1.** Immobilization of PbCDPS on biochar from polystyrene & wood waste, macroalgae, spent coffee and distillery waste, and in beads from alginate, alginate coated and chitosan. Rows in green highlight support systems with more than 30% of immobilization selected for the measurement of product formation (cHE).

| System | Immobilization (%) |
| --- | --- |
| *PbCDPS<sub>alg beads</sub>* | 100 |
| *PbCDPS<sub>alg beads</sub><sup>glut</sup>* | 100 |
| *PbCDPS<sub>alg/cot beads</sub>* | 100 |
| *PbCDPS<sub>alg/cot beads</sub><sup>glut</sup>* | 100 |
| *PbCDPS<sub>chi beads</sub>* | $9.72 \pm 2.61$ |
| ***PbCDPS*<sup>glut</sup><sub>chi beads</sub>** | 83.67 ± 4.66 |
| ***PbCDPS*<sub>biochar macroalgae</sub>** | 45.74 ± 3.65 |
| ***PbCDPS*<sup>glut</sup><sub>biochar macroalgae</sub>** | 29.65 ± 4.47 |
| ***PbCDPS*<sub>biochar spent coffee</sub>** | 30.25 ± 5.52 |
| ***PbCDPS*<sub>biochar spent coffee</sub> - CO<sub>2</sub>**<br>**activated** | 91.89 ± 4.11 |
| ***PbCDPS*<sup>glut</sup><sub>biochar spent coffee</sub>** | 36.17 ± 10.47 |
| ***PbCDPS*<sub>biochar distillery waste</sub>** | 17.03 ± 1.74 |
| ***PbCDPS*<sub>biochar distillery waste</sub> - CO<sub>2</sub>**<br>**activated** | <u>87.5 ± 0.98</u> |
| ***PbCDPS*<sup>glut</sup><sub>biochar distillery waste</sub>** | 33.08 ± 5.23 |
| ***PbCDPS*<sub>biochar poly & wood</sub>** | 27.22 ± 6.54 |
| ***PbCDPS*<sup>glut</sup><sub>biochar poly & wood</sub>** | 24.19 ± 3.25 |

FTIR measurements were performed using biochar samples to verify the chemical groups present at the material surface (see Fig. S1). FTIR spectra between 3600-3200 cm^-1^ reveal peaks corresponding to OH bonds of phenol or alcohol groups. However, the relatively small size of these peaks is due to the loss of moisture caused by the high temperatures reached in the gasification process^37^. The absorbance peak at 3000-2800 cm^-1^ represents the aliphatic C-H stretch vibration. They are poorly pronounced in all samples due to the degradation of aliphatic compounds at high gasification temperatures, resulting in a higher peak in mild pyrolysis temperatures. The absorbance peaks at 800 and 1600 cm^-1^ are attributed to the aromatic C-H stretch and the aromatic C=C stretch, respectively. In all biochar samples, peaks attributed to cellulose and hemicellulose (3200-3000 cm^-1^ for OH and 3100-3000 cm^-1^ for CH) are absent, due to the hemicellulose and cellulose being completely thermally degraded in biochar^38^. Double bonds could be due to the condensate aromatic structure observed in graphene. Representative peaks for C-H stretching (750-900 and 3050-3000 cm^-1^), C=C (1380-1450 cm^-1^), C-C and C-O stretching (1580-1700 cm-^1^) are present^39^. Bands between 1800 and 1500 cm^-1^ can be attributed to the C=O bond stretch of carboxylic acids and ketones. Fig. S1e, S1f and S1g show the FTIR spectra for beads. The broad peak at 3367 cm^-1^ was assigned to the free hydroxyl groups. Characteristic peaks of alginate were 1606 cm^-1^ and 1425 cm^-1^ for the C=O bond. The characteristics peaks of chitosan were at 1664 cm^-1^ for the amide I and 1544 cm^-1^ for the amide II^24^.

From scanning electron microscopy (SEM) studies, the biochar samples have different structural characteristics due to the different biomass morphology as shown in Fig. 2. The majority of the waste analyzed has a porous fiber structure typical of biomass except for the spent coffee (Fig. 2c and 3d), where the structure was not as well defined as in other samples, and deeper cavities suitable for the adsorption of molecules were observed. This result agrees with the pore size study (Table S1), as biochar from spent coffee was the material with the highest pore size. Biochar from macroalgae and distillery waste (Fig. 2a and 2e) have a honeycomb pore structure. Biochar from polystyrene & wood waste (Fig. 2h) have fewer pores produced due to the complex lignin cellulose structure typical of the wood. Additionally, images show that in general biochar has a high degree of macroporosity, which could permit a suitable adsorption of the protein and an effective interaction between enzymes and substrates^39^.

**Figure 2.**
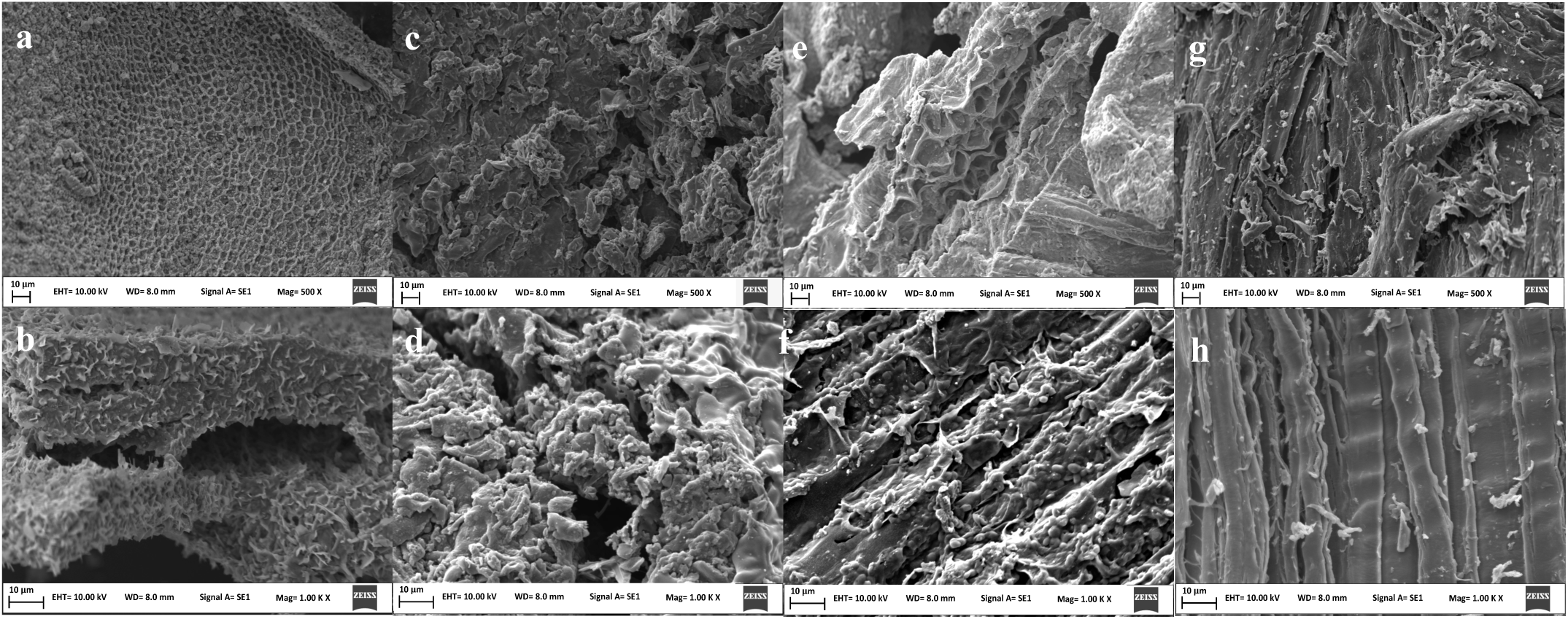
SEM images of biochar derived from (a) macroalgae at 500 x magnification, (b) macroalgae at 1000 x magnification, (c) spent coffee at 500 x magnification, (d) spent coffee at 1000 x magnification, (e) distillery waste at 500 x magnification, (f) distillery waste at 1000 x magnification, (g) polystyrene & wood waste at 500 x magnification and (h) polystyrene & wood waste at 1000 x magnification. All the samples were pyrolyzed at 550 °C., were gold-coated, and 10 kV were used for all the analysis.

**Figure 3.**
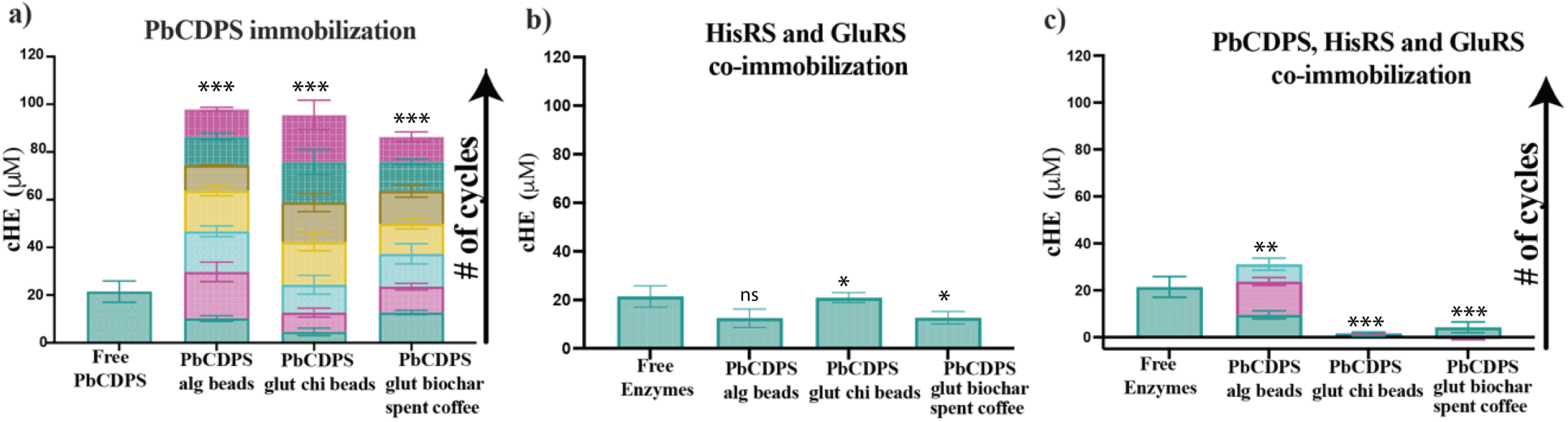
(a) Comparison of with free PbCDPS (not re-usable), with the reusability of PbCDPS immobilized in Ca-alginate beads by entrapment (*PbCDPS_alg beads_*), chitosan beads by covalent bonding 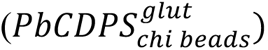, and biochar from spent coffee by covalent bonding 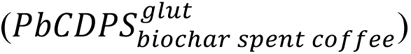. Same abbreviations were used for other panels in this figure. (b) Comparison of reusability for the co-immobilization of PbCDPS, HisRS and GluRS in different supports. (c) Co-immobilization of PbCDPS, HisRS and GluRS in different supports. Each colour in the stacked bar graph represents an additional cycle for the cHE production. All experiments were performed in triplicate, data are reported average ± SEM. One way ANOVA and Tukey’s test were performed using Graphpad Prism, and comparisons between free PbCDPS and other conditions are shown as follows: ns for not-statistically significantly different, * for p ≤ 0.05, ** for p ≤0.02 and *** for p ≤0.001.

### Immobilization of PbCDPS

#### Ca-alginate beads

PbCDPS was immobilized in alginate beads by the entrapment technique. With this approach, no PbCDPS was detected in the supernatant after the immobilization in all systems under comparison (Table 1), suggesting this support was a promising candidate for the activity assay of PbCDPS. Alginate is commonly used due to its stability, non-toxicity, and low cost. Multiple enzymes have been immobilized using this technique^33,40–42^. Previously, horseradish peroxidase (HRP) was immobilized in Ca-alginate beads and a maximum immobilization of 89 ± 5% was reported^43^, considerable but likely lower to what was achieved with *PbCDPS_alg beads_* and 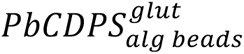. Two approaches for immobilization were employed, with and without functionalization, and in both cases no protein was detected after immobilization (Table 1). Immobilization of HRP in Ca-alginate beads using glutaraldehyde was achieved with an immobilization of 87%^44^.

#### Alginate-coated chitosan beads

Two approaches for immobilization were employed, with and without functionalization, and in both cases no protein was detected after immobilization (Table 1). High immobilization percentage has been reported with this system, such as 75% immobilization achieved for acrylamidase in chitosan coated alginate beads^34^. Immobilization of HRP in Ca-alginate beads using glutaraldehyde was achieved with an immobilization of 87%^44^. Reports indicate the mechanical strength is higher in coated beads than in simple alginate beads^34^. Additionally, the free hydroxyl groups in chitosan can react with other groups, which could explain the high levels of immobilization obtained.

#### Chitosan beads

Chitosan beads activated with glutaraldehyde provided a biocompatible support surface leading to a maximum immobilization of 83.27% (Table 1). This result is in agreement with previous reports for this support^45^. In this study, manganese peroxidase was immobilized onto glutaraldehyde active chitosan beads and 81.4% of immobilization was achieved. Glutaraldehyde functionalization is an important step for enzyme immobilization, as during the reaction between glutaraldehyde with chitosan generation of aldehyde groups on the bead surface could lead to side reactions with amino groups of the enzyme^46^. Additionally, it has been reported that chitosan beads could improve their mechanical resistance after functionalization due to the crosslinking of the polymeric chains of chitosan. In the present work, in the absence of glutaraldehyde only 9.72% of the enzyme was immobilized, and therefore the enzyme is likely mainly attached to the functional groups that glutaraldehyde provides and not to the support itself.

#### Biochar

In preliminary experiments (Table S2), the glutaraldehyde concentration and immobilization time were established. The immobilization of the PbCDPS on the biochar systems ranged between 17.03% and 45.74%. Comparing the immobilization time of chitosan beads (8 hours) with the biochar (0.5 hours), it is apparent that in a short time frame high loadings were not achieved, probably since PbCDPS will be mostly immobilized on the surface of beads, whereas more surface area is available with biochar and immobilization is completed with less time. Although PbCDPS was immobilized in the biochar systems, the precise mechanism underlying adsorption is unknown. However, we hypothesize that the enzyme is supported in the biochar by van der Walls forces. Multiple enzymes have been successfully immobilized using this approach on different materials^32,47,48^. For example, pepsin was immobilized on biochar obtained from pupuhna palm by adsorption and covalent bonding. A high immobilization efficiency (>95%) was obtained, possibly due to the porosity and pore diameter of the biochar and low molecular weight (35 kDa) of the enzyme^49^.

### Immobilized PbCDPS retains activity

To determine whether immobilized PbCDPS maintained activity after the immobilization process, a cut off for immobilization efficiency was set to >30% (Fig. 1, panel 3 on the “Prioritization of the different immobilization strategies used for PbCDPS”), and the activity assay for PbCDPS was performed only for the selected systems marked green on Table 2. PbCDPS trapped on alginate beads (*PbCDPS_alg beads_*) exhibits greater catalytic potential compared to other beads systems under evaluation, as this system had the highest concentration of cHE product after reaction. The high content of cHE produced with *PbCDPS_alg beads_* could be due to high porosity of alginate beads allowing necessary interactions between the PbCDPS and the aa-tRNA substrates^50^. Aa-tRNAs are large molecules (average molecular weight ∼25 kDa), and therefore production of cHE was highly affected by the porosity of the beads. However, adding glutaraldehyde to the immobilization or coating alginate beads with chitosan resulted in decreased product production. This could be due to glutaraldehyde acting as a crosslinking agent, as it was previously reported to provide a reinforcement to alginate beads through cross-linking within the matrix^51^. Glutaraldehyde could also lead to protein crosslinking and interfere with active site availability.

**Table 2.** cHE production from PbCDPS immobilized using different strategies. Rows in green are immobilization systems showing the highest concentration of cHE production, and further taken up for reusability testing.

| System | Immobilized<br>PbCDPS in<br>the reaction<br>(μg) | Immobilized<br>PbCDPS in<br>the reaction<br>(μM) | cHE (μM) | Ratio cHE<br>μM/μM of<br>PbCDPS<br>immobilized |
| --- | --- | --- | --- | --- |
| *PbCDPS<sub>alg beads</sub>* | 11.95 | 8.50 | 10.22<br>± 0.17 | 1.20 |
| *PbCDPS<sub>alg beads</sub><sup>glut</sup>* | 11.95 | 8.50 | 6.13 ±<br>0.93 | 0.72 |
| *PbCDPS<sub>alg:chi beads</sub><sup>glut</sup>* | 11.95 | 8.50 | 4.18 ±<br>0.27 | 0.49 |
| *PbCDPS<sub>chi beads</sub><sup>glut</sup>* | 14.6 | 10.38 | 8.72 ±<br>1.91 | 0.84 |
| *PbCDPS<sub>biochar macroalgae</sub>* | 14.6 | 10.38 | 7.23 ±<br>3.10 | 0.69 |
| *PbCDPS<sub>biochar macroalgae</sub><sup>glut</sup>* | 14.6 | 10.38 | 8.03 ±<br>1.65 | 0.77 |
| *PbCDPS<sub>biochar spent coffee</sub>* | 14.6 | 10.38 | 9.04 ±<br>0.08 | 0.87 |
| *PbCDPS<sub>biochar spent coffee-CO<sub>2</sub></sub>*<br>*activated* | 44.71 | 15.90 | ND | -- |
| *PbCDPS<sub>biochar spent coffee</sub><sup>glut</sup>* | 14.6 | 10.38 | 12.48 ±<br>1.06 | 1.20 |
| *PbCDPS<sub>biochar distillery waste</sub><sup>glut</sup>* | 14.6 | 10.38 | 8.69± 2.36 | 0.83 |
| *PbCDPS<sub>biochar distillery waste-CO<sub>2</sub></sub>*<br>*activated* | 42.58 | 15.14 | ND | -- |

ND: not determined as HPLC UV peak area is indistinguishable from noise. -- denotes unavailable data since immobilization was unsuccessful.

In the case of 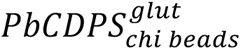, chitosan would be positively charged under the conditions employed while tRNAs would be negatively charged, increasing the likelihood of electrostatic attraction between tRNA and the support. Both glutaraldehyde concentration and biochar surface area could permit the immobilization of PbCDPS on the surface, not inside the material, and this immobilization could facilitate interactions between the enzymes and substrates participating in cHE production.

In the activity assays, a cascade reaction produces the final product cHE. If the aaRS enzymes (HisRS and GluRS) are free in solution they will generate aa-tRNA products, which then must act as substrates for the entrapped PbCDPS enzyme. This reaction was confirmed by HPLC-MS, since cHE will only be observed if the cascade reaction is completed successfully (Fig. S2). *PbCDPS_alg beads_* system was improved by varying the concentration of PbCDPS in beads. Fig. S3a shows that the maximum ratio of cHE/PbCDPS concentration was achieved with 13.9 µg of PbCDPS, as larger amounts of immobilized PbCDPS did not translate to increased cHE formed. Similarly, in the case of the immobilization on biochar from spent coffee (Fig. S3(b)), different loadings of PbCDPS were tested, and the maximum activity was obtained when 15 µg was employed. In the case of 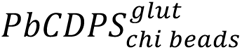, the immobilization percentage was similar for all the loadings tested. Diffusional obstacles may be expected using larger enzyme loadings, and that could reduce the observed activity of immobilized protein^52^. We hypothesize this could be due to the large size of aa-tRNAs and the fact that aaRS enzymes are not embedded in the support, and therefore aa-tRNAs would have to diffuse in and out of the support, decreasing the amount of product generated.

To test this hypothesis and determine whether the tRNA pool employed remains free in solution or is adsorbed to beads during and after the reaction in the three selected supports, an RNA denaturing gel was run with the supernatant following immobilization (see Fig. S4). After reaction, the tRNA pool concentration was higher in the supernatant than in the support. This could be due to the large molecular weight of tRNA, requiring large pores to penetrate into beads or biochar. Additionally, because tRNA was mostly free in solution, other proteins in the cascade most likely need to remain free or at least solvent accessible in order to complete the sequence of reactions leading to cHE. Only in the chitosan beads was the concentration of tRNA pool similar in the beads and in the supernatant, likely due to electrostatic interactions between tRNA and chitosan. Additionally, protein gel electrophoresis was employed to test whether aaRS enzymes were free in solution or adsorbed to beads during and after the reaction. Fig. S5 shows that HisRS and GluRS were present mainly in the supernatant of all systems. Consequently, a successful support cannot hinder contact between substrates and enzymes, and therefore the physical properties of the support employed will play an important role in the selection of an ideal support.

### Reusability of immobilized PbCDPS

Reusability of immobilized PbCDPS was studied for the selected systems in batch reactions 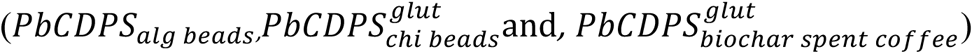. The number of cycles in which immobilized PbCDPS retained activity is a marker for immobilization success. In *PbCDPS_alg beads_*, the amount of cHE detected increased as cycles progressed (Fig. 3a). This could be due to product entrapped in the support after each reaction cycle, so a quenching experiment was performed (Fig. S6) comparing product recovered after the system was completely quenched and when only supernatant, free of beads, was recovered. Product yield was higher when system was completely quenched until the fourth cycle. This could be due to bead saturation with product.

A similar experiment evaluating cHE product entrapment in the support was carried out for chitosan beads and biochar (Fig. S6). The amount of product after the reaction was similar when the supernatant or entire system were analyzed. This was confirmed in the reusability experiments with biochar and chitosan (Fig. 3a). In a similar study, Bilal et al. ^35^ studied the reusability of manganese peroxidase immobilized onto chitosan beads, observing that immobilized enzymes retained up to 5 cycles of their initial activity. Whereas Santos et al.^49^ reported an efficiency of 85% after seven cycles in the immobilization of pepsin on biochar. Immobilized laccase on biochar from wood maintained up to 80% of its initial activity after five cycles.

### Co-immobilization of PbCDPS and aaRS enzymes

The production of cyclic dipeptides involves a cascade reaction between three enzymes - two aaRS and PbCDPS. We therefore proceeded to immobilize the three enzymes on the same system. Firstly, the two aaRS were immobilized on the selected supports (alginate beads, chitosan beads and biochar from spent coffee). The activity of the cHE was detected in the three systems when HisRS and GluRS were immobilized and PbCDPS was free in the reaction mixture, demonstrating immobilized aaRS enzymes retain activity following immobilization (Fig. 3b). Then, the immobilization of the three enzymes (PbCDPS, HisRS and GluRS) involved in the cascade reaction for the cyclic dipeptide production was carried out (Fig. 3c). With alginate beads, the amount of cHE product was similar to when PbCDPS was immobilized alone. In the case of biochar from spent coffee and the chitosan beads, cHE was obtained at lower amounts than when only PbCDPS was immobilized (Fig. 3c). Alginate beads have been used successfully employed for the co-immobilization and recycling of other enzymes. Li et al.^52^ reported that in porous materials, the co-immobilization could be troublesome, since some enzymes could be localized in the micropores and others on the surface of the material, causing mass transfer issues between the substrates and enzyme catalysts taking part in the reaction.

When the reusability of the co-immobilized enzymes was tested (Fig. 3c), alginate beads were the most promising option for immobilization. However, after the third reaction cycle enzymatic activity showed signs of decline. In this case, as previously observed, some cHE was trapped inside the beads after the first cycle. The findings presented in Fig. S5 suggest that the decline in activity over the cycles could be attributed to challenges in mass transfer to and from the beads, rather than the loss of enzyme activity within the reaction. This could be because tRNA molecules remain outside the support structure during and after the reaction, which could present challenges to active site access for the enzymes involved in the cascade.

## Conclusions and significance

In this work, we established the immobilization of PbCDPS leading to the production of the cyclodipeptide cHE. PbCDPS is involved in the biosynthesis of cyclic dipeptide natural products with pharmaceutical applications, requiring two large aminoacyl-tRNAs as substrates. Although immobilization has been used extensively, application to enzymes involved in the biosynthesis of natural products are scarce. When free in solution, PbCDPS is an unstable enzyme, catalyzing few turnovers before denaturation. Immobilization improved enzyme stability, as after seven reaction cycles the enzyme maintained catalytic turnover. Furthermore, we demonstrated the co-immobilization of PbCDPS and other enzymes participating in the cascade reaction to produce cyclic dipeptides (HisRS, GluRS) can be successfully performed. tRNA synthetases are widely employed in transcription/translation commercial kits and have extensive application in molecular and chemical biology. Therefore, our work demonstrates the feasibility of enzyme immobilization, alone or in cascade, when large complex substrates such as tRNAs are required. While the successful immobilization of PbCDPS represents a significant advancement in stabilizing enzymes for cyclic dipeptide synthesis, further work to optimize its efficiency is needed, including enzyme engineering. Exploring the scalability of this process for industrial applications holds immense promise, as ∼65 % drugs approved from 1981 to 2019 are either natural products or molecules derived from natural products^53^. Advancing this approach toward industrial-scale implementation could offer a pathway for efficient, sustainable, and large-scale synthesis of cyclic dipeptide natural products with diverse pharmaceutical applications.

## Author Contributions

L.A.R. performed experiments on enzyme immobilization and activity and writing original draft, E.S contributed to PbCDPS purification and tRNA pool generation, C.M.C., R.P.S. and A.D.B. planned the project and participated in data analysis and interpretation. E.M.M.M., R.P.S, A.D.B. and C.M.C reviewed and edited the manuscript.

All authors have given approval to the final version of the manuscript.

## Acknowledgements

CMC and ES are funded by the Wellcome trust (217078/Z/19/Z), LAR is funded by Consejo Nacional de Humanidades Ciencia y Tecnología (CONAHCYT) Mexico (486638). RPS and EMMM are funded by the project: Development of smart edible coating for the preservation of berries I025-IAMSM005-C3-T1-T, of the Challenge-Based Research Funding program of the Tecnologico de Monterrey.

## Supporting Information

The Supporting Information is available and contains FTIR spectra for different supports employed for immobilization, standard curve for cHE quantification, details on biochar particle size and immobilization efficiency, gels showing location of proteins and tRNA during the PbCDPS catalyzed reaction, and effect on different quenching strategies in calculating cHE yield.

## Synopsis

We immobilized three enzymes in a biocatalytic cascade to produce cyclodipeptide natural products. Our strategy increases recyclability and yield with challenging and complex tRNA substrates.

