## Supporting information for "Overcoming the challenges of cascade reactions and complex substrates in natural product biocatalysis: Immobilization of a cyclodipeptide synthase"

**Figure S4. RNA denaturing gel. The identification of the lanes are as follows: 1-molecular weight marker; 2-supernatant after reaction with** $PbCDPSbiochar\ spent\ coffeeglut$**; 3-supernatant after reaction with** $PbCDPSalg\ beads$**; 4-supernatant after reaction with** $PbCDPSchi\ beadsglut$**; 5-**$PbCDPSbiochar\ spent\ coffeeglut$ **support from; 6-**$PbCDPSalg\ beads$**; support after reaction; 7-**$PbCDPSchi\ beadsglut$**support after the reaction; 8-molecular weight marker; 9-tRNA pool.** .......... 5

**Figure S5. SDS-PAGE for immobilization mixture loaded with 20 µM total protein per lane. The identification of the lanes are as follows: 1- molecular weight ladder; 2-PbCDPS; 3- HisRS; 4-GluRS; 5- supernatant after reaction with** $PbCDPSchi\ beadsglut$**; 6-supernatant after reaction with** $PbCDPSalg\ beads$**; 7-supernantat after reaction with** $PbCDPSbiochar\ spent\ coffeeglut$**; 8-**$PbCDPSchi\ beadsglut$**support after reaction; 9-**$PbCDPSalg\ beads$ **support after reaction; 10- support from** $PbCDPSbiochar\ spent\ coffeeglut$. .......... 6

#### Supporting Figures

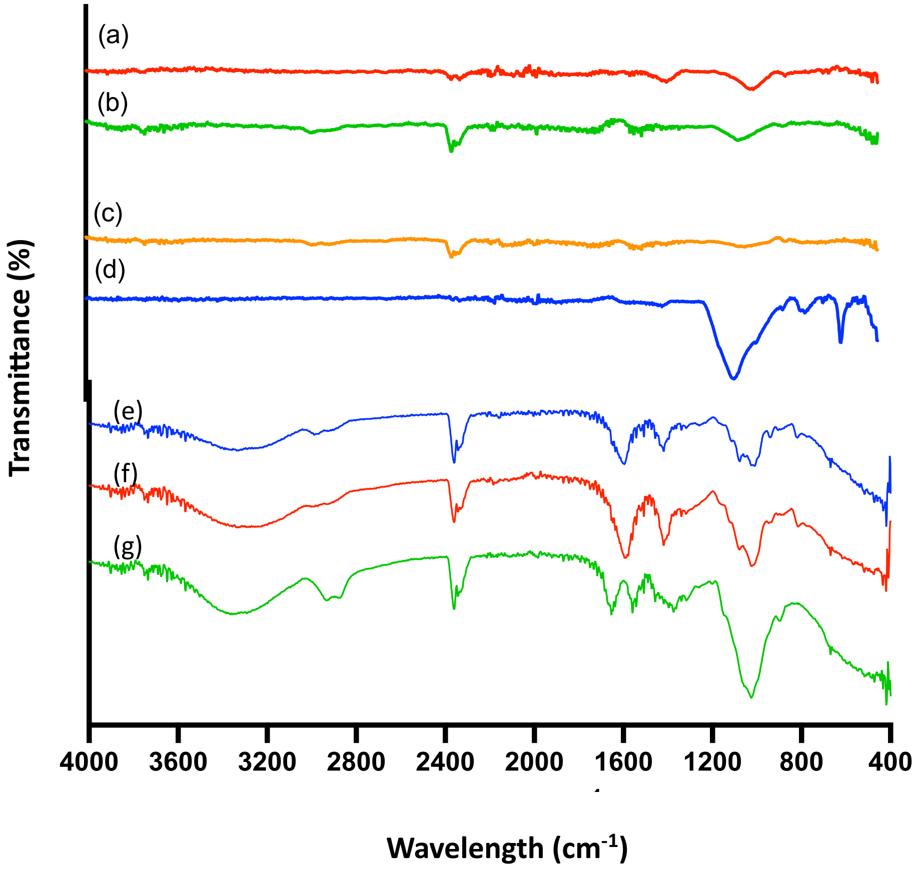

Figure S1. FTIR spectra of biochar samples from spent coffee (a), distillery waste (b), plastic & wood waste (c), macroalgae (d), Ca-alginate beads €, alginate coated beads (f), chitosan beads (g).

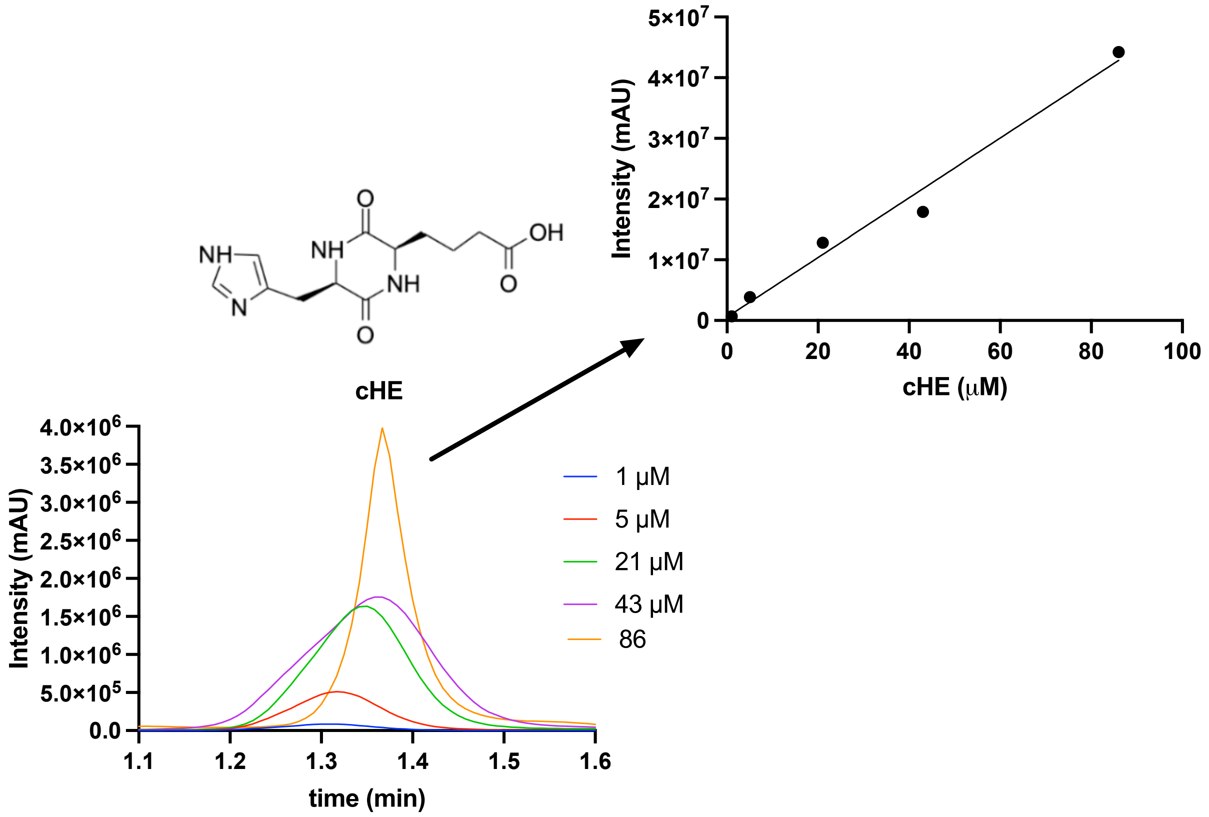

Figure S2. HPLC chromatogram of the purification of cHE at 214 nm and calibration curve used a for the quantification of the cHE. Structure of the cHE was showed.

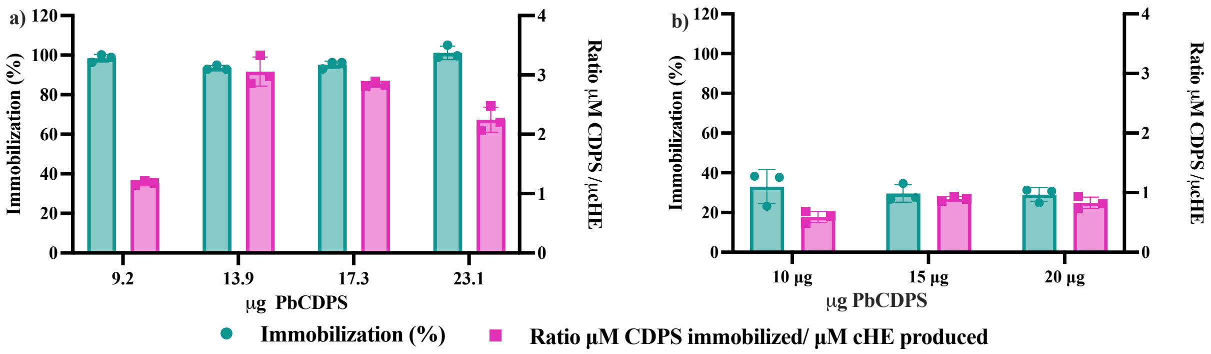

Figure S3. (a) Effect of PbCDPS loading in the immobilization in alginate beads, and ratio of PbCDPS/cHE). (b) Effect of PbCDPS loading in the immobilization on biochar from spent coffee, and ratio of PbCDPS/cHE.

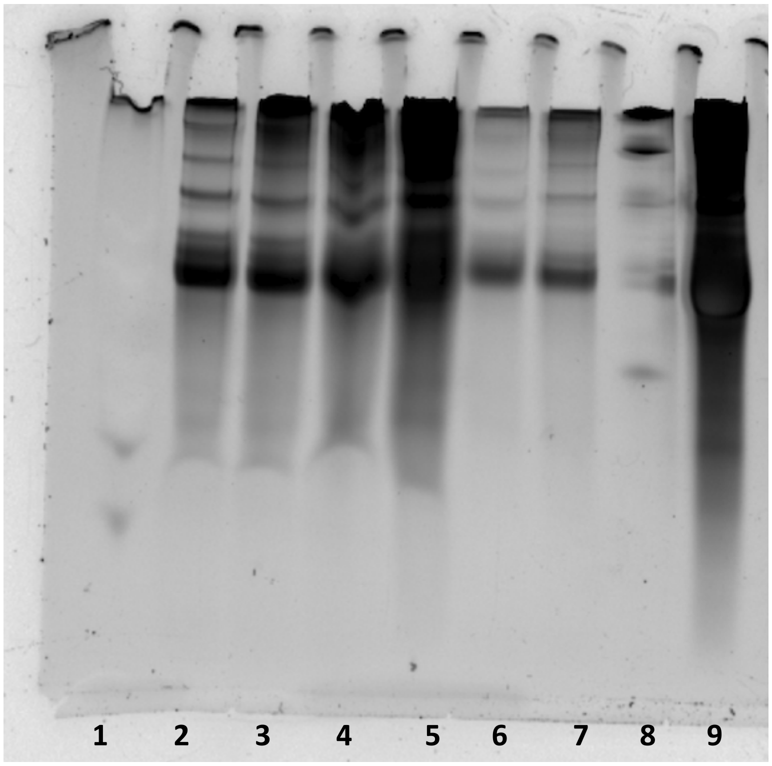

Figure S4. RNA denaturing gel. The identification of the lanes are as follows: 1-molecular weight marker; 2-supernatant after reaction with $PbCDPS_{biochar\ spent\ coffee}^{glut}$; 3-supernatant after reaction with $PbCDPS_{alg\ beads}$; 4-supernatant after reaction with $PbCDPS_{chi\ beads}^{glut}$; 5-$PbCDPS_{biochar\ spent\ coffee}^{glut}$ support from; 6-$PbCDPS_{alg\ beads}$; support after reaction; 7- $PbCDPS_{chi\ beads}^{glut}$support after the reaction; 8-molecular weight marker; 9-tRNA pool.

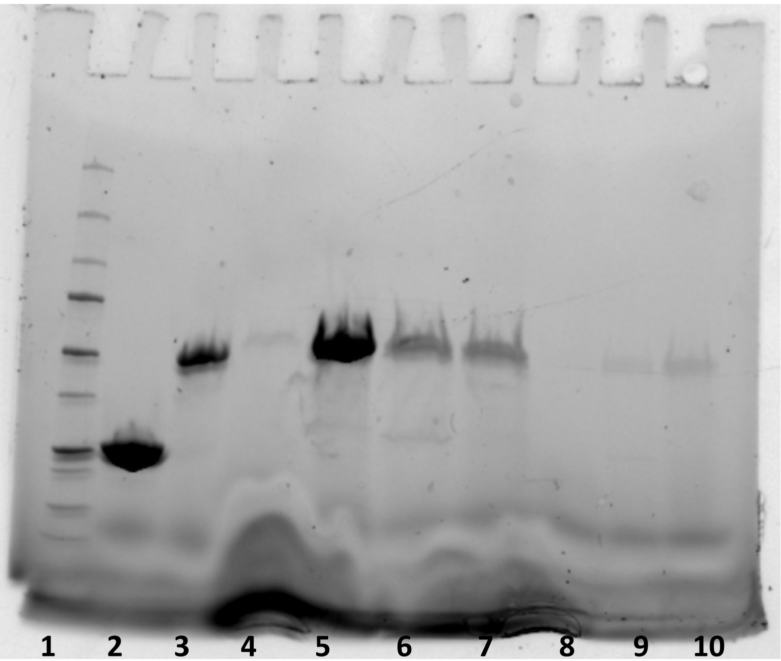

Figure S5. SDS-PAGE for immobilization mixture loaded with 20 $\mu$M total protein per lane. The identification of the lanes are as follows: 1- molecular weight ladder; 2- PbCDPS; 3- HisRS; 4-GluRS; 5- supernatant after reaction with $PbCDPS^{glut}_{chi\ beads}$; 6- supernatant after reaction with $PbCDPS_{alg\ beads}$; 7-supernantat after reaction with $PbCDPS^{glut}_{biochar\ spent\ coffee}$; 8-$PbCDPS^{glut}_{chi\ beads}$support after reaction; 9- $PbCDPS_{alg\ beads}$ support after reaction; 10- support from $PbCDPS^{glut}_{biochar\ spent\ coffee}$.

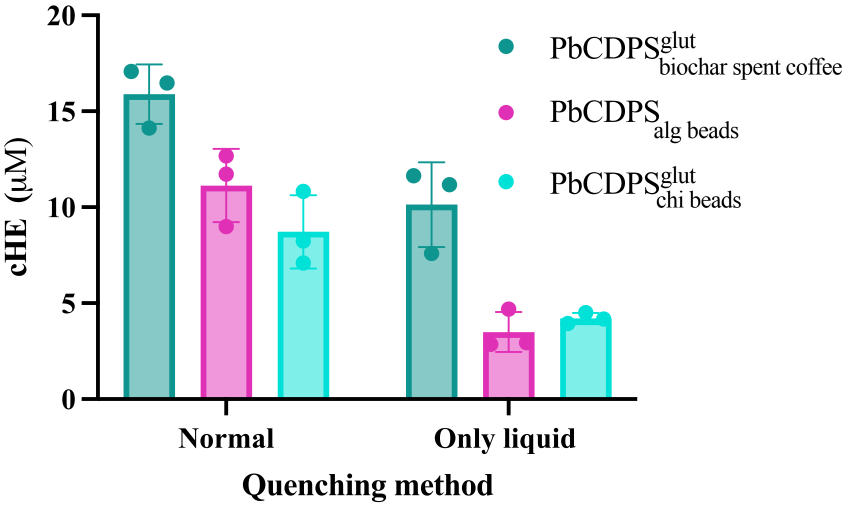

Figure S6. Effect of different quenching strategies for the measurement of cHE the selected supports ($\mathrm{PbCDPS_{alg\ beads}}$,$\mathrm{PbCDPS^{glut}_{chi\ beads}}$and, $\mathrm{PbCDPS^{glut}_{biochar\ spent\ coffee}}$)

### Supporting Tables

Table S1. BET specific surface area, pore volume and pore size of biochar from macroalgae, spent coffee, distillery waste and plastic & wood waste with and without activation with $CO_2$.

| **Support** | **Surface area ($m^2/g$)** | **Pore volume (cc/g)** | **Pore size (nm)** |
|---|---|---|---|
| Macroalgae | 73.09 | 0.092 | 1.641 |
| Macroalgae $CO_2$ | 46.92 | 0.126 | 0.017 |
| Spent coffee | 3.738 | 0.005 | 3.791 |
| Spent coffee $CO_2$ | 14.41 | 0.026 | 0.014 |
| Distillery waste | 6.793 | 0.011 | 2.769 |
| Plastic & wood waste | 17.81 | 0.003 | 2.769 |
| Plastic & wood waste $CO_2$ | 18.11 | 0.3867 | 0.0101 |

Table S2. Immobilization of PbCDPS on activated carbon and cHE product formation.

| **Time (h)** | **Glutaraldehyde concentration (%)** | **Immobilization (%)** | **cHE (mM)** |
|---|---|---|---|
| 0.5 | | 23.5 | 2.72 |
| 1 | 1 | 22.2 | ND |
| 20 | | 48.9 | 3.43 |
| 0.5 | | 14.8 | 5.23 |
| 1 | 2 | 14 | ND |
| 20 | | 45.8 | 7.59 |

ND: not determined as HPLC UV peak area is indistinguishable from noise.

Table S3. Loading effect of PbCDPS on the immobilization of chitosan activated beads.

| **PbCDPS (μg)** | **Immobilization (%)** |
|---|---|
| 7.3 | 84.58 ± 3.95 |
| 14.6 | 83.67 ± 4.66 |
| 29.2 | 87.44 ± 1.97 |
| 58.4 | 86.87 ± 1.78 |

Table S4. Co-immobilization of HisRS and Glu-RS in alginate beads, chitosan beads activated with glutaraldehyde and biochar from spent coffee activated with glutaraldehyde.

| **System** | **Immobilization (%)** |
|---|---|
| $PbCDPS_{alg\ beads}$ | 100 ± 1.15 |
| $PbCDPS^{glut}_{chi\ beads}$ | 52.48 ± 9.10 |
| $PbCDPS^{glut}_{biochar\ spent\ coffee}$ | 24.47 ± 7.99 |
